# GeneDMRs: an R package for Gene-based Differentially Methylated Regions analysis

**DOI:** 10.1101/2020.04.11.037168

**Authors:** Xiao Wang, Dan Hao, Haja N. Kadarmideen

**Author notes:** **Corresponding authors**, Correspondence to Xiao Wang or Haja N. Kadarmideen.

## Abstract

DNA methylation in gene or promoter or gene body could restrict/promote the gene transcription. Moreover, methylation in the gene regions along with CpG island regions could modulate the transcription to undetectable gene expression levels. Therefore, it is necessary to investigate the methylation levels within the gene, gene body, CpG island regions and their overlapped regions and then identify the gene-based differentially methylated regions (GeneDMRs). Here, R package *GeneDMRs* aims to facilitate computing gene based methylation rate using next generation sequencing (NGS)-based methylome data. The user-friendly R package *GeneDMRs* is presented to analyze the methylation levels in each gene/promoter/exon/intron/CpG island/CpG island shore or each overlapped region (e.g., gene-CpG island/promoter-CpG island/exon-CpG island/intron-CpG island/gene-CpG island shore/promoter-CpG island shore/exon-CpG island shore/intron-CpG island shore). Here, we used the public reduced representation bisulfite sequencing (RRBS) data of mouse (GSE62392) for evaluating software and found novel biologically significant results to supplement the previous research. The R package *GeneDMRs* can facilitate computing gene based methylation rate to interpret complex interplay between methylation levels and gene expression differences or similarities across physiological conditions or disease states.

## 1. Introduction

Generally, gene expression is restricted by DNA methylation. However, the presence of methylation in gene or promoter or gene body could result in promotion of gene transcription. Irizarry et al. (2009) revealed the correlation between substantial portion of DNA methylation sites and gene expression along the genome. DNA methylation in promoters usually restricts the genes in a long-term stabilization of repressed states, while most gene bodies are also extensively methylated in different status; therefore, methylation of such regions can be the potential therapeutic targets (Jones, 2012; Yang et al., 2014). CpG islands, regions of high density of DNA methylation of cytosine and guanine dinucleotides (CpGs), are playing an important role in gene regulation and transcriptional repression (Goldberg et al., 2007). Moreover, the shore regions beyond CpG islands are also involved in modulating gene expression (Irry et al., 2009; Doi et al., 2009).

Identifying causal relationships via genotype–phenotype correlations is a substantial challenge and many studies across life sciences try to integrate multi-omics datasets in that effort (Suravajhala et al., 2016). Recently, one of the largest genetic study investigated global gene expression and DNA methylation patterns in 265 human skeletal muscle biopsies from the FUSION study with > 7 million genetic variants. This integrated approach led to potential causal mechanisms for eight physiological traits: height, waist, weight, waist–hip ratio, body mass index, fasting serum insulin, fasting plasma glucose, and type 2 diabetes (Taylor et al., 2019). This underlines the importance of having gene-based methylation rates to integrate with differential expression or co-expression across physiological and phenotypic or disease states.

Studying DNA methylation patterns in biological samples using next generation sequencing (NGS) methods are becoming increasingly common. There are several tools available to detect differentially methylated cytosine (DMC) (e.g., R package *IMA* (Wang et al., 2012), *MethylKit* (Akalin et al., 2012)) or differentially methylated region (DMR) (e.g., R package *ELMER* (Silva et al., 2018), *MethylMix* (Gevaert, 2015; Cedoz et al., 2018),*Minfi* (Aryee et al., 2014), *MIRA* (Lawson et al., 2018), *RnBeads* (Assenov et al., 2014; Müller et al., 2019)). These packages mainly focus on the specific differentially methylated regions like genes (DMGs) from cancer dataset (Gevaert, 2015; Cedoz et al., 2018) or only promoters (DMPs) (Assenov et al., 2014; Müller et al., 2019). However, detections of DMR based on gene body features associated with CpG islands are scarce, such as the DMRs in all exons (DMEs) and introns (DMIs) or a specific exon and intron. To the best of our knowledge, there are no tools that detect the DMP/DME/DMI/DMG associated with CpG islands/CpG island shores. We present here a user-friendly R package *GeneDMRs* (https://github.com/xiaowangCN/GeneDMRs) to facilitate computing gene based methylation rate using next generation sequencing (NGS) based methylome data. *GeneDMRs* analyzes the methylation levels in each gene/promoter/exon/intron/CpG island/CpG island shore or each overlapped region (e.g., gene/promoter/exon/intron CpG island and gene/promoter/exon/intron CpG island shore). We evaluated the R package *GeneDMRs* using the publicly available reduced representation bisulfite sequencing (RRBS) data from mouse (Accession ID: GSE62392).

## 2. Materials and Methods

### 2.1 Data structure in DNA methylation

Genome-wide DNA methylation analysis are mainly based on three approaches, i.e., enzyme digestion, affinity enrichment and bisulfite conversion (Laird, 2010). Whole genome bisulfite sequencing (WGBS) aims to find the whole methylome (Frommer et al., 1992) while reduced representation bisulfite sequencing (RRBS) primarily focuses on the enrichment of CpG-rich regions by recognizing the site CmCGG after restriction enzyme *MspI* digestion (Meissner et al., 2005), but both techniques rely on bisulfite conversion. From WGBS or RRBS data, cytosine site information (e.g. chromosome and position) and its methylation status can be obtained using available bioinformatics tools. *GeneDMRs* package can directly use the resulting methylation *coverage* file (i.e., *.bismark.cov*) from *Bismark* software or similar file with chromosome, start position, end position, methylation percentage, number of methylated read and number of unmethylated read (five or six columns). With such dataset, we provide below the statistical framework to compute gene-based methylation rate.

### 2.2 Gene-based DMRs and analysis workflow

The gene-based regions could be divided into single window, gene, promoter, exon, intron, CpG island and CpG island shore and their overlapped feature regions including gene-CpG island, gene-CpG island shore, promoter-CpG island, promoter-CpG island shore, exon-CpG island, exon-CpG island shore, intron-CpG island and intron-CpG island shore (Figure 1).

**Figure 1.**
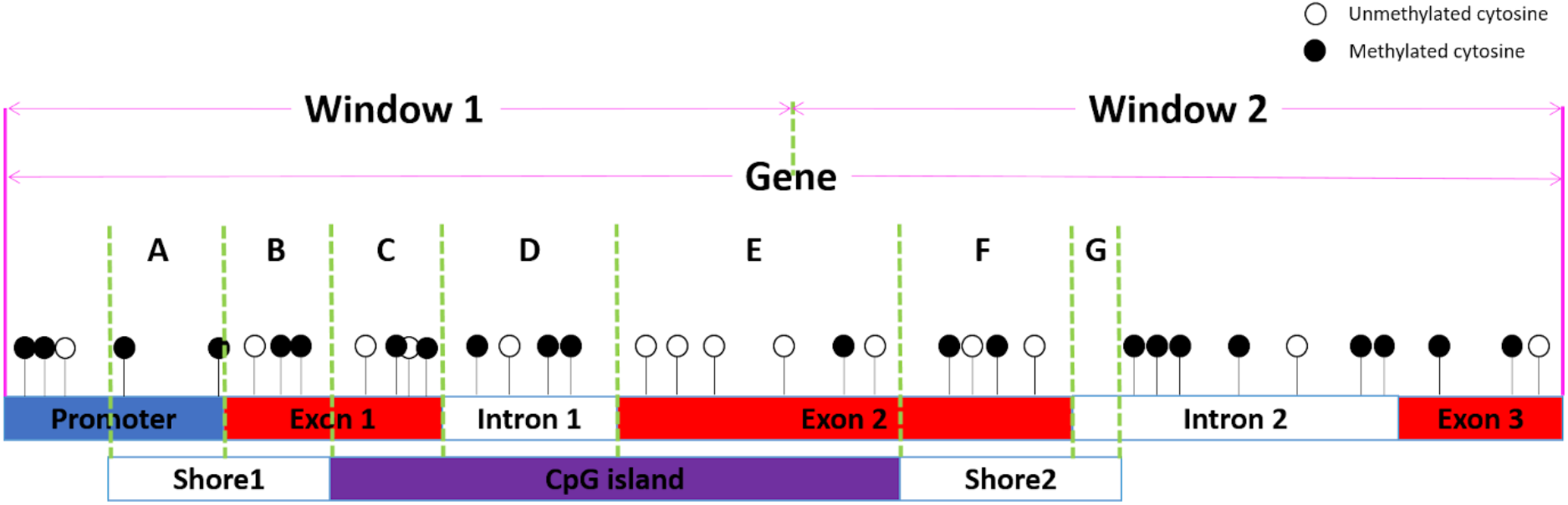
The analyzed targets in the *GeneDMRs* package including widows, genes (promoters, exons, introns), CpG islands (CpGis, Shores) and the overlapped feature regions (e.g., **A**: Promoter-Shore1, **B**: Exon1-Shore1, **C**: Exon1-CpGi, **D**: Intron1-CpGi, **E**: Exon2-CpGi, **F**: Exon2-Shore2, **A** + **B**: Gene-Shore1, **C** + **D** + **E**: Gene-CpGi, **F** + **G**: Gene-Shore2).

The methylation mean of a cytosine site by weighting for one group (a case or control) is calculated by (1):

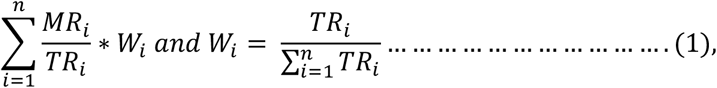

where *MR_i_* and *TR_i_* are the methylated and total reads number at a given cytosine site of individual *i*, *n* is the total number of individuals in one group and *W_i_* is the weight of reads of individual *i*.

For a window/gene (promoter, exon, intron)/CpGi/other overlapped region (Figure 1) of one group, the methylation mean by weighting is calculated by (2):

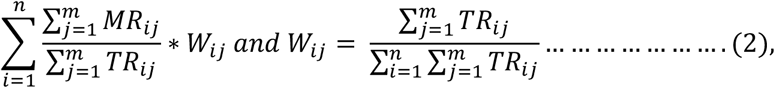

where *MR_ij_* and *TR_ij_* are the methylated and total reads number of the involved cytosine/DMC site *j* at a given gene/CpGi/other region of individual *i*, *m* is the total number of cytosine/DMC sites involved in this region, *n* is the total individual number of one group and *W_ij_* is the weight of reads of the involved cytosine/DMC site *j* of individual *i*. For the target region, the cytosine/DMC within the region is selected, and then methylation mean of each group is calculated. Here, the DMC sites refer to the differentially methylated cytosine sites after `Significant_filter(siteall_Qvalue, qvalue = 0.01, methdiff = 0.05)`. Thus, if the users want to use the DMC sites for the methylation mean, they should calculate the *Q*-values and methylation difference by `Logic_regression()` and filter out the DMCs by `Significant_filter()` at first (Figure 2). This step was also used in our previous study for methylation difference calculation to discover hyper and hypo-methylated DMGs (Wang and Kadarmideen, 2019a).

**Figure 2.**
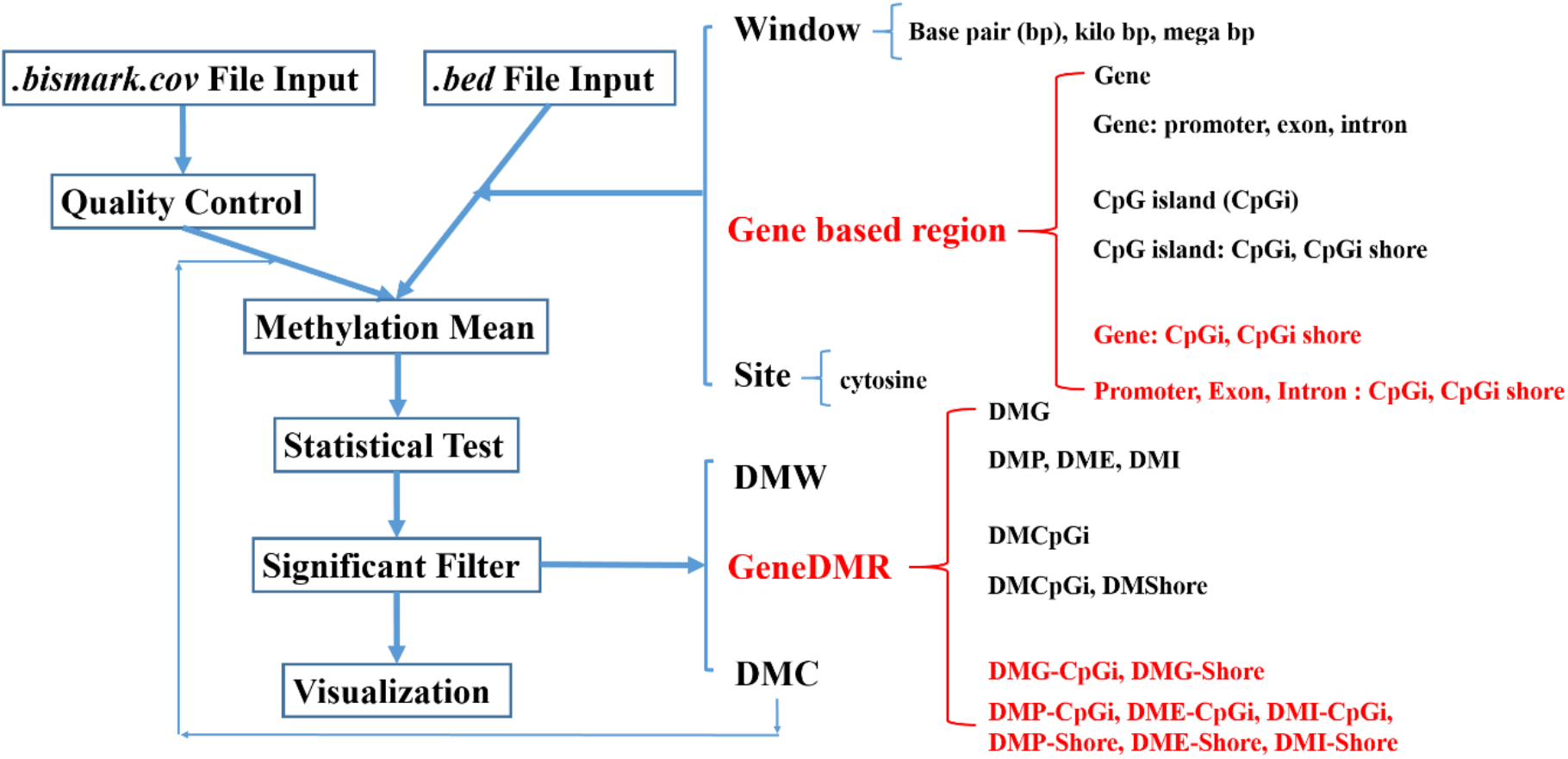
Overall workflow of *GeneDMRs* package.

Logistic regression model were used to fit methylation levels with the different groups following R package *MethylKit* (Akalin et al., 2012):

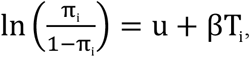

where π_i_ is the methylation mean at a given window or gene-based region or site, u is the intercept, and T_i_ is the group indicator.

More categorical variables can also be incorporated in this model as the additional covariates by `Logic_regression(covariates = NULL)`. Chi-squared (χ2) test was used to determine the statistical significance of methylation differences among different groups and then to achieve the *P*-values. To account for multiple hypothesis testing, *P*-values can be adjusted to *Q*-values by various methods, e.g., “bonferroni”, “holm” (Holm, 1979), “hochberg” (Hochberg, 1988), “hommel” (Hommel, 1988), “BH” (Hochberg, 1995), “fdr” (Hochberg, 1995) and “BY” (Benjamini and Yekutieli, 2001).

Differentially methylated windows (DMWs) or gene-based DMRs or DMCs (Figure 2) are mainly filtered by *Q*-values and methylation level differences between two groups, e.g., `Significant_filter(qvalue = 0.01, methdiff = 0.05)`. The methylation difference can be calculated in `Logic_regression(diffgroup = c(“group1”, “group2”))` by selecting any two groups. The differentially methylated genes (DMGs) will be defined as the hyper/hypo-methylated gene when the methylation difference is positive/negative after case-control comparison (e.g., group2 - group1).

Based on gene-based regions, DMRs for specific regions could be detected, such as genes (DMGs), promoters (DMPs), exons (DMEs), introns (DMIs), CpG islands (DMCpGis), CpG island shores (DMShores) and the overlapped regions like gene-CpG islands (DMG-CpGis), gene-CpG island shores (DMG-Shores), promoter-CpG islands (DMP-CpGis), promoter-CpG island shores (DMP-Shores), exon-CpG islands (DME-CpGis), exon-CpG island shores (DME-Shores), intron-CpG islands (DMI-CpGis) and intron-CpG island shores (DMI-Shores) (Figure 2). In addition, the ordinal positions of exons and introns were identified for each gene, which can be used in the further analysis, for example the correlations of methylation levels between all promoters and all first exons. The overall workflow of *GeneDMRs* package includes file input, quality control, methylation mean calculation, statistical test, significant filter and results visualization (Figure 2).

### 2.3 Application to real data

The reduced representation bisulfite sequencing (RRBS) data for testing the developed method was download from Gene Expression Omnibus (GEO) with the accession number GSE62392 (https://www.ncbi.nlm.nih.gov/geo/query/acc.cgi?acc=GSE62392). The downloaded RRBS data was originally generated from RRBS of sorted common myeloid progenitor (CMP) populations that were isolated from 3 pools of G0 as control group and 2 pools of G5 as case group of mice samples (Colla et al., 2015). Mouse data used here is an example and *GeneDMRs* package is applicable to all species. We applied several pre- and post-mapping quality control (QC) to this data as follows. Adapters and reads less than 20 bases long of RRBS data were trimmed by *Trimmomatic* software (version 0.36) (Bolger et al., 2014). The clean reads were mapped to the mice reference genome by *Bowtie 2* software (version 2.3.3.1) (Langmead and Salzberg, 2012). The house mouse (*Mus musculus*) reference genome (mm10) used in this study was downloaded from the University of California Santa Cruz (UCSC) website (http://hgdownload.soe.ucsc.edu/goldenPath/mm10/bigZips/mm10.2bit). The *.2bit* file was subsequently converted to *.fasta* file by *twoBitToFa* software (http://hgdownload.cse.ucsc.edu/admin/exe/linux.x86_64/twoBitToFa). Finally, read coverages of detected methylated or unmethylated cytosine sites and their methylation percentages were obtained by using *Bismark* software (version 0.19.0) (Krueger and Andrews, 2011). In this study, we only considered the numbers of methylated and unmethylated cytosines in CpG sites for each sample and the non-CpG (CHG and CHH, H representing A/C/T) sites were discarded.

### 2.4 Input and quality control

The resulting methylation *coverage* files from *Bismark* software of five mouse RRBS data were directly used as input in *GeneDMRs* package. The public mouse (mm10) *bed* file (i.e., *.bed*) for Reference Sequence (refseq) and CpG island (cpgi) database were downloaded from the UCSC website (http://genome.ucsc.edu/cgi-bin/hgTables). RefSeq and CpG island *bed* files were used as input files in *GeneDMRs* package which then can output four files (inputrefseqfile, inputcpgifile, inputgenebodyfile and inputcpgifeaturefile) by altering the *feature* parameter in the reading function, e.g., `Bedfile_read(feature = TRUE/FALSE). Bedfile_read()` function divides each gene of refseq *bed* file into four gene body features (i.e., promoters, exons, introns and TSSes) and each CpG island of cpgi *bed* file into two CpG island features (i.e., CpG islands and CpG island shores) based on R package *genomation* (Akalin et al., 2015). Moreover, `Bedfile_read()` function annotates specific gene to each promoter by the further step. If the strand of one promoter is “+”/“-”, the middle position of this promoter will be the start/end position of the matched specific gene. However, for the specific genes with more than one National Center for Biotechnology Information (NCBI) ID, *GeneDMRs* package will choose the first one. For example, the adenosine A1 receptor gene (Adora1) has four NCBI IDs (i.e., NM_001291930, NM_001282945, NM_001039510 and NM_001008533) and only the first ID (NM_001291930) will be chosen.

When a polymerase chain reaction (PCR) experiment suffers from duplication bias, some clonal reads will impair accurate determination of methylation (Akalin et al., 2012). In addition, lower read coverages (e.g., lower than 10) will cause the biases for methylation percentage calculation (Wang and Kadarmideen, 2019b). Therefore, cytosines with a percentile of read coverage higher than the 99.9^th^ and read coverages lower than 10 were discarded for the qualified reads by `Methfile_QC(high_quantile = 99.9, low_coveragenum = 10)`.

### 2.5 Biological enrichment for the differentially methylated genes (DMGs)

After Significant_filter() function for DMGs, these genes with methylation differences can be used for biological enrichment. The enrichments for GO terms and pathways are analyzed and visualized by `Enrich_plot(enrichterm = c(“GO”, “pathway”), category = TRUE/FALSE)` based on R package *clusterProfiler* (Yu et al., 2012). If the `category = TRUE`, the enrichment will display in hyper-methylated and hypo-methylated categories. In addition, the differentially expressed genes (DEGs) with Log fold change (LogFC) information can also be used in `Enrich_plot(expressionfile_significant = NULL)`, then the visualized enrichment will be in four categories that are hyper/hypo-methylated and up/down-regulated genes. The up/down-regulated DEG can be defined when the LogFC is positive/negative or derived from the previous research results. Here, we use the previous results for multiple-category enrichments that are downregulated and upregulated genes in G4/G5 compared to G0 CMP (fdr = 0.05) of mice samples (https://ars.els-cdn.com/content/image/1-s2.0-S1535610815001403-mmc2.xlsx) (Colla et al., 2015).

## 3. Results and Discussion

### 3.1 Comparisons of different R packages for methylation analysis

Currently, a series of R packages can analyze methylation data to detect DMCs or DMRs (Table 1). Most of them are not however completely focusing on the regions in genes or within gene bodies or CpG islands and hence *GeneDMRs* package could be a complementary tool to obtain methylation levels at these levels. As shown in Table 1, *ELMER v.2* package reconstructs altered gene regulatory network (GRN) by combining enhancer methylation and gene expression (Silva et al., 2018). *IMA* (Wang et al., 2012) and *MethylKit* (Akalin et al., 2012) aim at genome-wide cytosine sites analysis for BeadChip and RRBS data, respectively. Generally, *methyAnalysis*, *MethylationArrayAnalysis* and *Minf* are packages for specific purposes, where *methyAnalysis* applies CpG island information to visualize in the heatmap plot and *Minfi* can find the hypomethylation blocks (Aryee et al., 2014). If considering methylated genes, *MethylMix* package mainly focuses on identifying disease specific hypo and hypermethylated genes and it defines the methylation difference of a methylation state with the normal methylation state (Gevaert, 2015; Cedoz et al., 2018), while *RnBeads* package could consider the gene, gene promoter, CpG island and genomic tiling regions [15, 16]. Overall, none of these R packages works for gene components, but *GeneDMRs* package is extended to exon and intron regions, and their interactions with CpG island features. In addition, the performance of was tested in the personal computer (CPU: 2.70 GHz, RAM: 8.00 GB) comparing with *MethylKit* package (Akalin et al., 2012). For all the reference genes, *GeneDMRs* package takes around 15 minutes while gene body dataset interacted with CpG island dataset requires the longest time, thus, the performance of *GeneDMRs* package is generally related to the number of analyzed targets (Figure 3).

**Table 1.** Comparisons of different R packages for methylation analysis.

| R package | Target | Analysis feature | Issued time |
| --- | --- | --- | --- |
| *ELMER* v.2 (Silva et al., 2018) | DMR | Reconstruct altered gene regulatory network (GRN) by combining enhancer methylation and gene expression | 2018 |
| *IMA* (Wang et al., 2012) | Site-level and region-level methylation | Summarization for individual site as well as annotated region | 2012 |
| *methyAnalysis* | DMR | Chromosome location based DNA methylation analysis and heatmap plot with CpG island | 2018 |
| *MethylationArrayAnalysis* | Probe-wise differential methylation and DMR | Differential variability analysis, estimating cell type composition and gene ontology testing | 2019 |
| *MethylKit* (Akalin et al., 2012) | Base or region of DNA methylation | Functions for clustering, sample quality visualization, differential methylation analysis and annotation feature | 2012 |
| *MethylMix* (Gevaert, 2015)<br>*/MethylMix 2.0* (Cedoz et al., 2018) | DMR of gene | Automate the construction of DNA-methylation and gene expression dataset from The Cancer Genome Atlas (TCGA) | 2015/2018 |
| *Minfi* (Aryee et al., 2014) | Differentially methylated position (DMP) and DMR | Block finding to identify hypomethylation block | 2014 |
| *MIRA* (Lawson et al., 2018) | DMR | Take advantage of genome-scale DNA methylation data to assess regulatory activity | 2018 |
| *RnBeads* (Assenov et al., 2014)<br>*/RnBeads 2.0* (Müller et al., 2019) | DMR of gene/promoter/CpG island | DNA methylation-based prediction of age and sex; LOLA-based region set enrichment analysis for biological interpretation | 2014/2019 |

**Figure 3.**
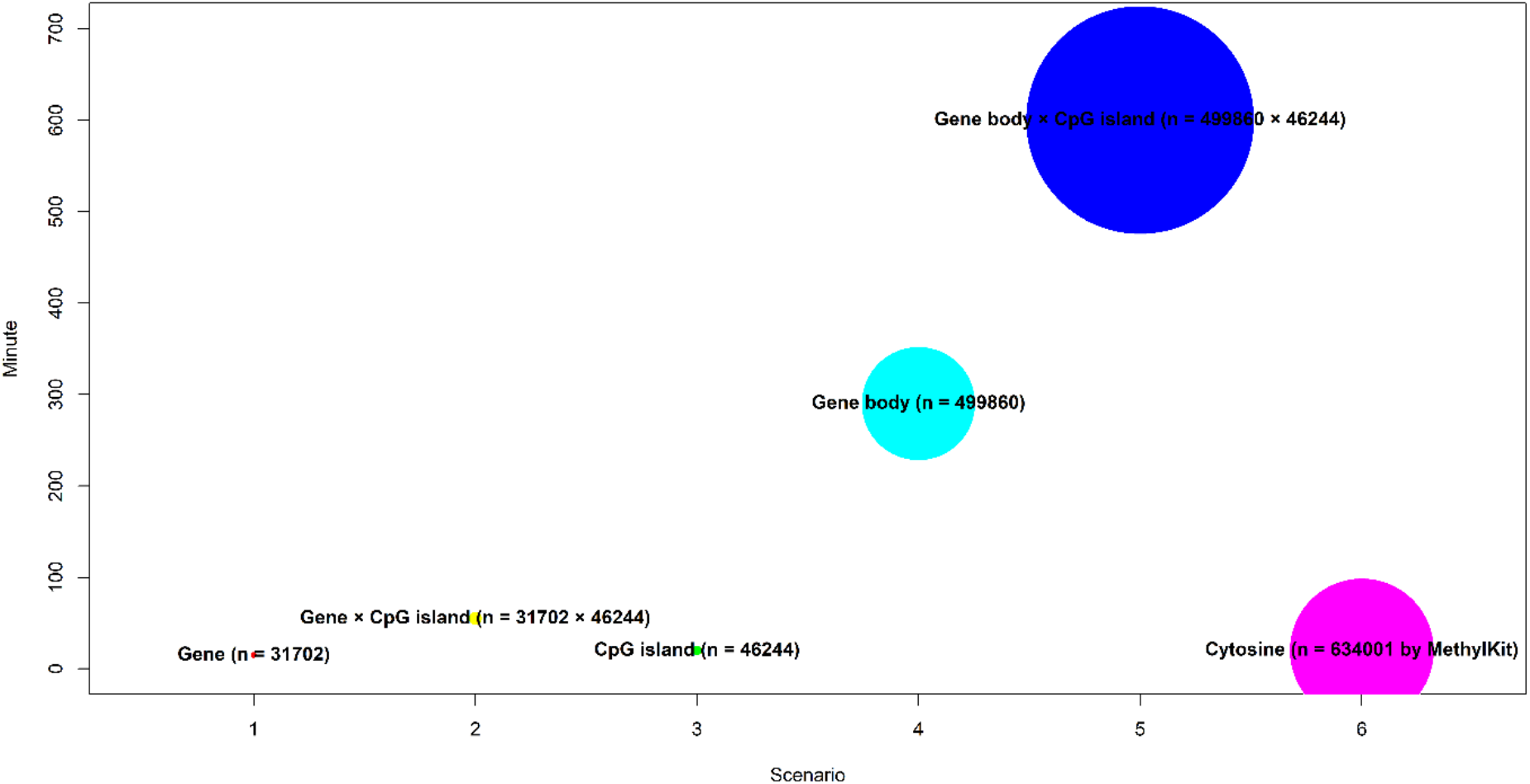
The performance of *GeneDMRs* package.

### 3.2 Differentially methylated gene-based regions and cytosine sites

In the final step, five methylation *coverage* files from *Bismark* software were used in *GeneDMRs* package and their statistical summary is listed in supplementary table 1. The *GeneDMRs* package will automatically read the files with the file name like “1_1”, “1_2” and “2_1” for group and replicate numbers. The methylation patterns of all genes and DMGs in different CpG island regions by `Group_cpgfeature_boxplot()` and `Genebody_cpgfeature_boxplot()` are shown in supplementary figure 1. Results suggest that the methylation levels of DMGs were higher than before and they are the same of CpG islands higher than shores (Supplementary figure 1). The all dataset for genes (`regiongeneall_Qvalue`), genes with CpG island features (`regiongeneall_cpgfeature_Qvalue`), gene bodies with CpG island features (`genefeatureall_cpgfeature_Qvalue`) and cytosine sites (`genefeatureall_cpgfeature_Qvalue`) after `Logic_regression()` are listed in Supplementary file 1, 2, 3 and 4, respectively.

The methylation difference of all the cytosine sites involved in the gene were centralized to a mean, so statistical power seemed be lower than before (Figure 4 and Supplementary figure 2). In addition, *GeneDMRs* package can detect different gene body regions (e.g., promoter, exon and intron), CpG island regions (e.g., CpGi and shore regions) and their overlapped regions by `Methmean_region(cpgifeaturefile = inputcpgifeaturefile/NULL, featureid = “ c(“chr1”,“chr2”)/all/alls”, featurename = c(“promoters”,“exons”,“introns”,“TSSes”)/c(“CpGisland”, “Shores”))` for different methylation mean calculations. According these results, we found that *DNMT3A* was a hypo-methylated (NM_001271753) gene but the gene and one intron interacted in both CpG island and shore features were in hyper-methylation status when G5 CMP was compared to G0 CMP (Supplementary file 1, 2 and 3). Therefore, *GeneDMRs* package can accurately find significantly and biologically methylated gene body and CpG island regions along the whole genome and supplement the previous research (Colla et al., 2015).

**Figure 4.**
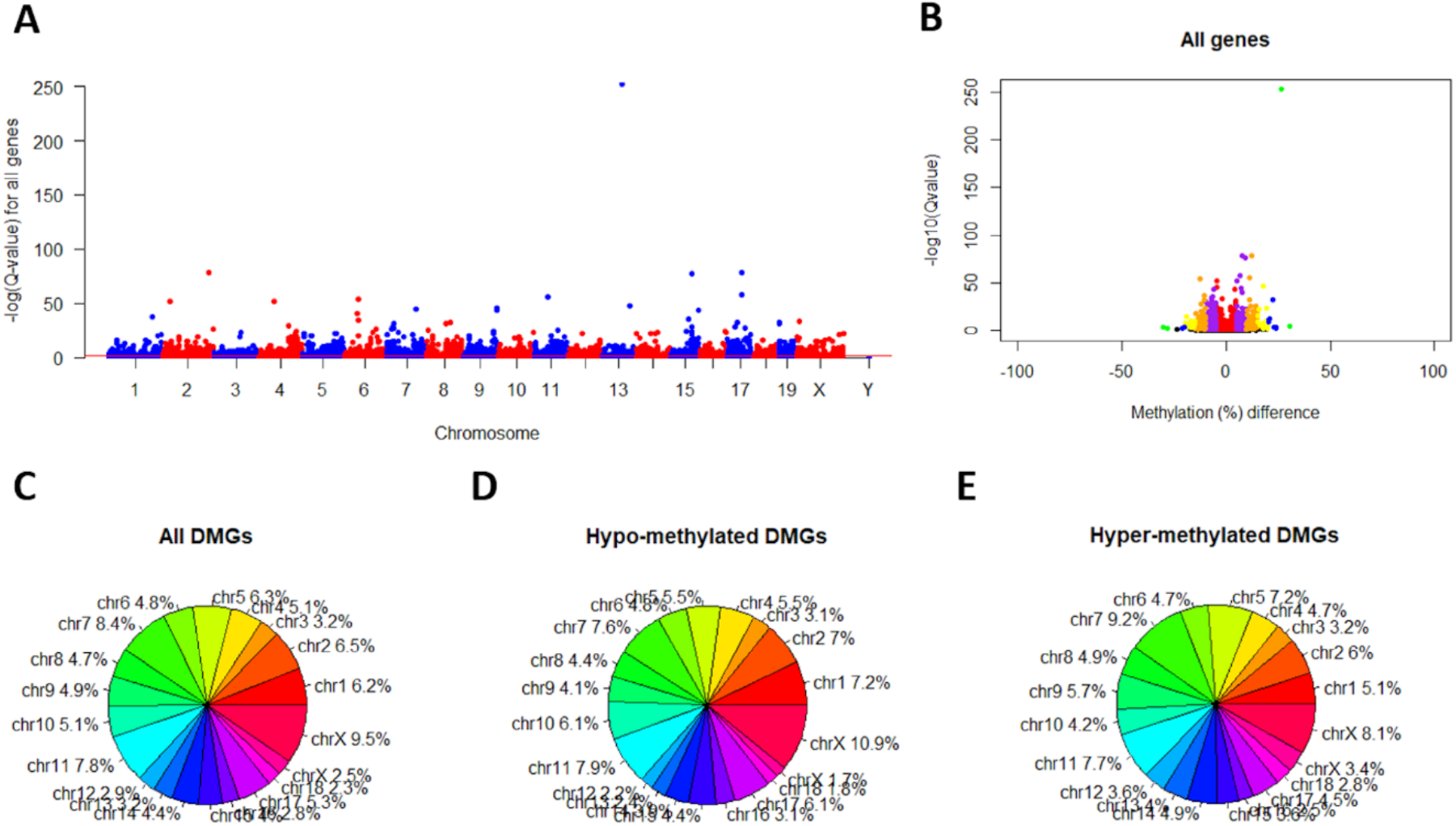
(**A**) Manhattan plots for all genes. Note: The red line indicates the significant level of Q-value < 0.01. (**B**) Methylation differences in all genes. Note: Plots showing red, purple, orange, yellow, blue and green colors indicate genes with a Q-value less than 0.01 and methylation difference (%) greater than 0, 5, 10, 15, 20 and 25, respectively. (**C**), (**D**) and (**E**) Percentages of all, hypo-methylated and hyper-methylated DMGs in different chromosomes, respectively.

If we only use the DMCs to recalculate the methylation mean by replacing the RRBS cytosine sites, i.e., `DMC_methfile_QC(inputmethfile_QC, siteall_significant)`, the methylation difference will be more obvious than before (Supplementary figure 3). The DMC-based methylation levels could represent the whole methylations for gene-based regions when the DMCs in one gene are involved in the important parts that affect the transcription. For WGBS data, statistical efficiency can be potentially improved by removing globally unmethylated sites with less methylation differences, because the total number of hypotheses affects the *Q*-values by the rank of combined *P*-values (Huh et al., 2017). The global DMC-based methylation levels (Figure 5) can be realized by `Circos_plot(inputcytofile, inputmethfile_QC, inputrefseqfile, inputcpgifeaturefile)` based R package *RCircos* (Zhang et al., 2013).

**Figure 5.**
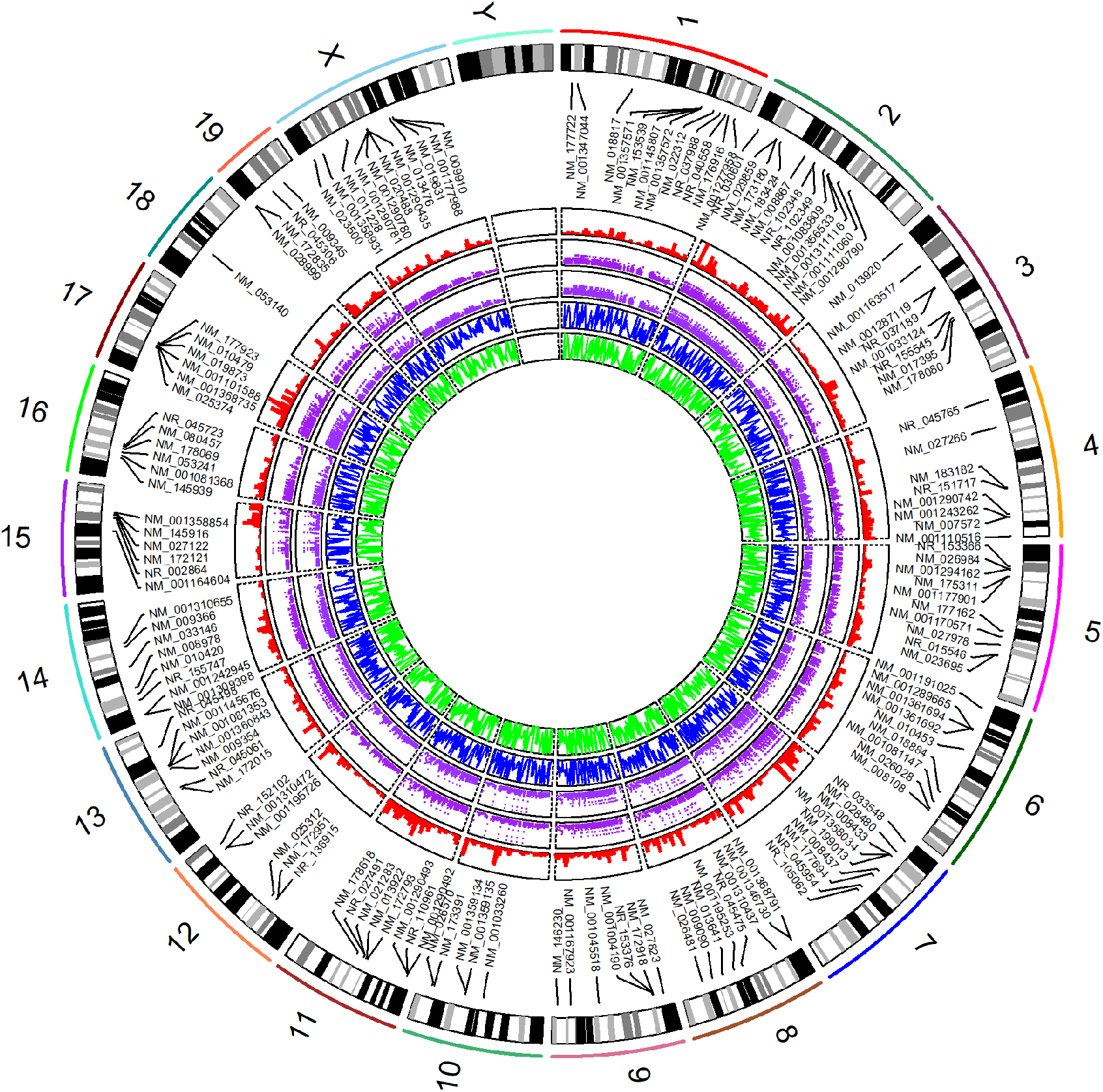
Circular graph of the global methylation levels. Note: From the outermost track to innermost circle, the circles indicate genome chromosomes (i.e., mouse), DMGs, gene densities, CpG island densities, CpG island shore densities and methylation levels. The densities and methylation levels were calculated by 1,000,000 base pair (bp) windows, i.e., `Window_divide(windowbp = 1000000)`.

### 3.3 Biological enrichment for DMGs

The enrichments for groups, GO terms and pathways can be analyzed and visualized with/without categories following R packages *clusterProfiler* (Yu et al., 2012). For example, the GO terms can be visualized in no/one/two categories (Figure 6) by incorporating hyper/hypo-methylated and up/down-regulated gene information. Thus, based on the DMGs and enrichments for GO term and pathway, *GeneDMRs* package can help to detect the specific significant regions, reveal the biological mechanism and enhance the previous studies that methylation pattern changes in specific-regions were involved in causing diseases (Colla et al., 2015).

**Figure 6.**
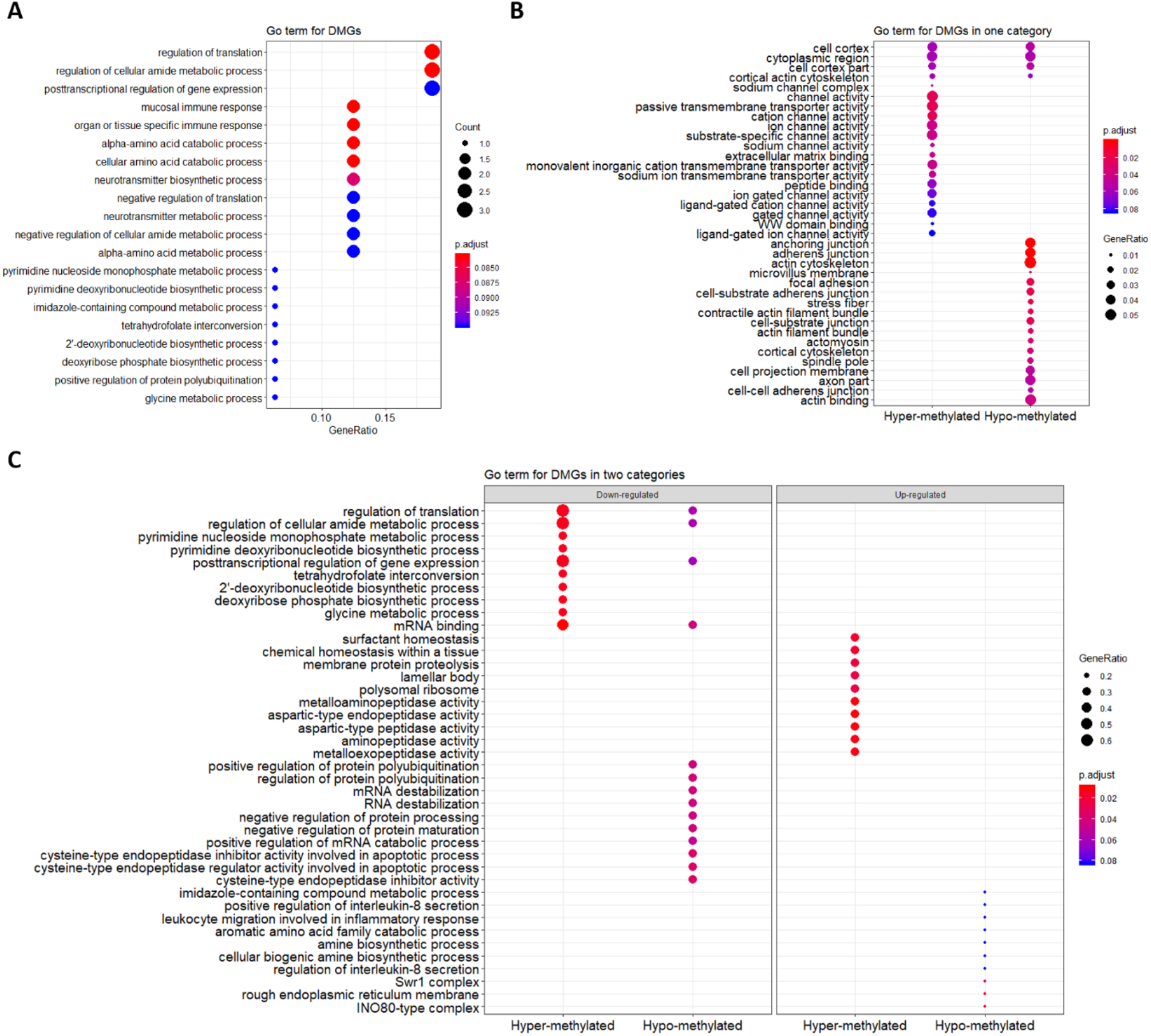
GO term enrichments. (**A**) GO terms without category. (**B**) GO terms with one category of hyper/hypo-methylated genes. (**C**) GO terms with two categories of hyper/hypo-methylated and up/down-regulated genes.

## 4. Summary

Currently, there is no easy-to-use R package that could compute methylation levels at the gene based level. *GeneDMRs*, a user-friendly R package, can facilitate computing gene based methylation rate using NGS-based methylome data. This package aims to analyze the methylation levels in gene/promoter/exon/intron/CpG island/CpG island shore and their overlapped regions. Then, the differentially hyper/hypo-methylated genes can be visualized for enrichments of GO terms and pathways and reveal the biological mechanism accordingly. Such gene-based methylation analyses contributes to interpreting complex interplay between methylation levels and gene expression differences or similarities across physiological conditions or disease states.

## List of abbreviations

Adora1: Adenosine A1 receptor gene
CMP: Common myeloid progenitor
CpG: Cytosine and guanine dinucleotide
DEG: Differentially expressed gene
DMC: Differentially methylated cytosine
DMCpGi: Differentially methylated CpG island
DME: Differentially methylated exon
DMG: Differentially methylated gene
DMI: Differentially methylated intron
DMP: Differentially methylated promoter
DMR: Differentially methylated region
DMShore: Differentially methylated CpG island shore
DMW: Differentially methylated window
GeneDMRs: Gene-based differentially methylated regions
GEO: Gene Expression Omnibus
GRN: Gene regulatory network
LogFC: Log fold change
NCBI: National Center for Biotechnology Information
NGS: Next generation sequencing
PCR: Polymerase chain reaction
QC: Quality control
RRBS: Reduced representation bisulfite sequencing
UCSC: University of California Santa Cruz
WGBS: Whole genome bisulfite sequencing

## Availability and Implementation

GeneDMRs is freely available at https://github.com/xiaowangCN/GeneDMRs

## Author information

### Affiliations

*Quantitative Genomics, Bioinformatics and Computational Biology Group, Department of Applied Mathematics and Computer Science, Technical University of Denmark, Kongens Lyngby, Denmark*

Xiao Wang & Haja N. Kadarmideen

*College of Animal Science and Technology, Northwest A&F University, China. Department of Molecular Biology and Genetics, Aarhus University, Denmark*.

Dan Hao

### Contributions

XW developed and implemented the method and *GeneDMRs* package, with supervision of HNK. DH gave feedback on package development and tested the final package. HNK interpreted the results from application of this package. XW wrote the manuscript. DH and HNK improved the manuscript. All authors read and approved the final manuscript.

## Supplementary materials

Supplementary table 1. Statistical summary of data source.

Supplementary figure 1. (**A**) Methylation patterns of all genes for different groups and gene bodies in different CpG island regions. (**B**) Methylation patterns of all DMGs for different groups and gene bodies in different CpG island regions. Note: P value is calculated by the methylation comparison between CpG island and CpG island shore with Student’s t-tests.

Supplementary figure 2. (**A**) Manhattan plots for all cytosine sites. Note: The red line indicates the significant level of Q-value < 0.01. (**B**) Methylation differences in all cytosine sites. Note: Plots showing red, purple, orange, yellow, blue and green colors indicate genes with a Q-value less than 0.01 and methylation difference (%) greater than 0, 5, 10, 15, 20 and 25, respectively. (**C**), (**D**) and (**E**) Percentages of all, hypo-methylated and hyper-methylated cytosine sites/DMCs in different chromosomes/gene bodies/CpG islands, respectively.

Supplementary figure 3. (**A**) Heat map cluster for methylation levels of all DMGs (n = 246). (**B**) Heat map cluster for methylation levels of all DMC-based DMGs (n = 2022). Note: DMGs and DMC-based DMGs were filter by `Significant_filter(qvalue = 0.01, methdiff = 0.1)`.

Supplementary file 1. Details of 20,837 genes with chromosomes, positions, methylation levels, read numbers, *P*-values, *Q*-values and methylation differences.

Supplementary file 2. Details of 14,822 genes interacted by CpG island features with chromosomes, positions, methylation levels, read numbers, *P*-values, *Q*-values and methylation differences.

Supplementary file 3. Details of 41,562 gene bodies interacted by CpG island features with chromosomes, positions, methylation levels, read numbers, *P*-values, *Q*-values and methylation differences.

Supplementary file 4. Details of 634,001 cytosines with chromosomes, positions, methylation levels, read numbers, *P*-values, *Q*-values and methylation differences.

