## Supplementary File for "GeneDMRs: an R package for Gene-based Differentially Methylated Regions analysis"

Supplementary table 1. Statistical summary of data source.

| Sample | Name in *GeneDMRs* package | Clean read pair | Uniquely mapping efficiency | Total cytosine number | Cytosine methylation rate in CpG context | Cytosine methylation rate in CHG context | Cytosine methylation rate in CHH context |
| --- | --- | --- | --- | --- | --- | --- | --- |
| G0-CMP1 | 1_1 | 30,015,396 | 62.1% | 201,134,947 | 21.6% | 0.3% | 0.3% |
| G0-CMP2 | 1_2 | 30,843,564 | 63.0% | 205,241,712 | 24.4% | 0.4% | 0.3% |
| G0-CMP3 | 1_3 | 32,558,450 | 62.4% | 215,559,294 | 23.1% | 0.4% | 0.3% |
| G5-CMP1 | 2_1 | 28,780,302 | 61.4% | 193,333,818 | 19.9% | 0.3% | 0.3% |
| G5-CMP2 | 2_2 | 31,798,819 | 60.7% | 204,493,356 | 24.9% | 0.3% | 0.3% |

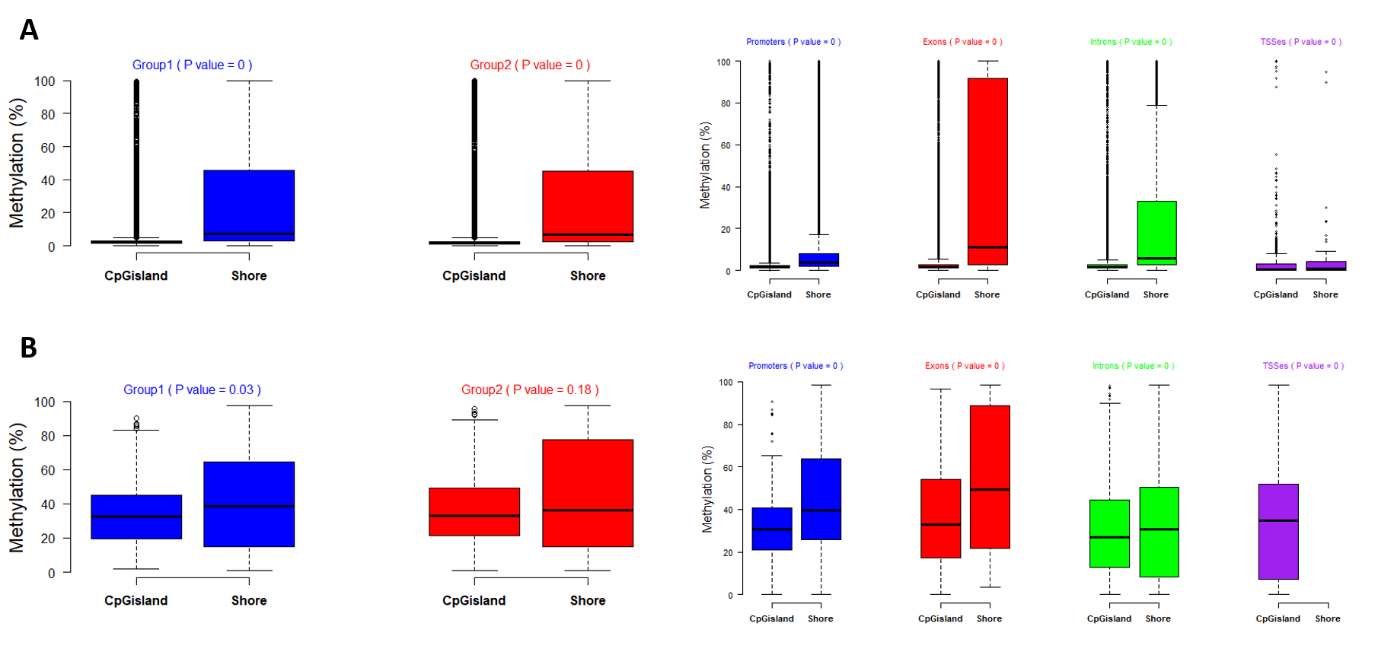

Supplementary figure 1. (**A**) Methylation patterns of all genes/cytosine sites for different groups and gene bodies in different CpG island regions. (**B**) Methylation patterns of all DMGs/DMCs for different groups and gene bodies in different CpG island regions. Note: P value is calculated by the methylation comparison between CpG island and CpG island shore with Student’s t-tests.

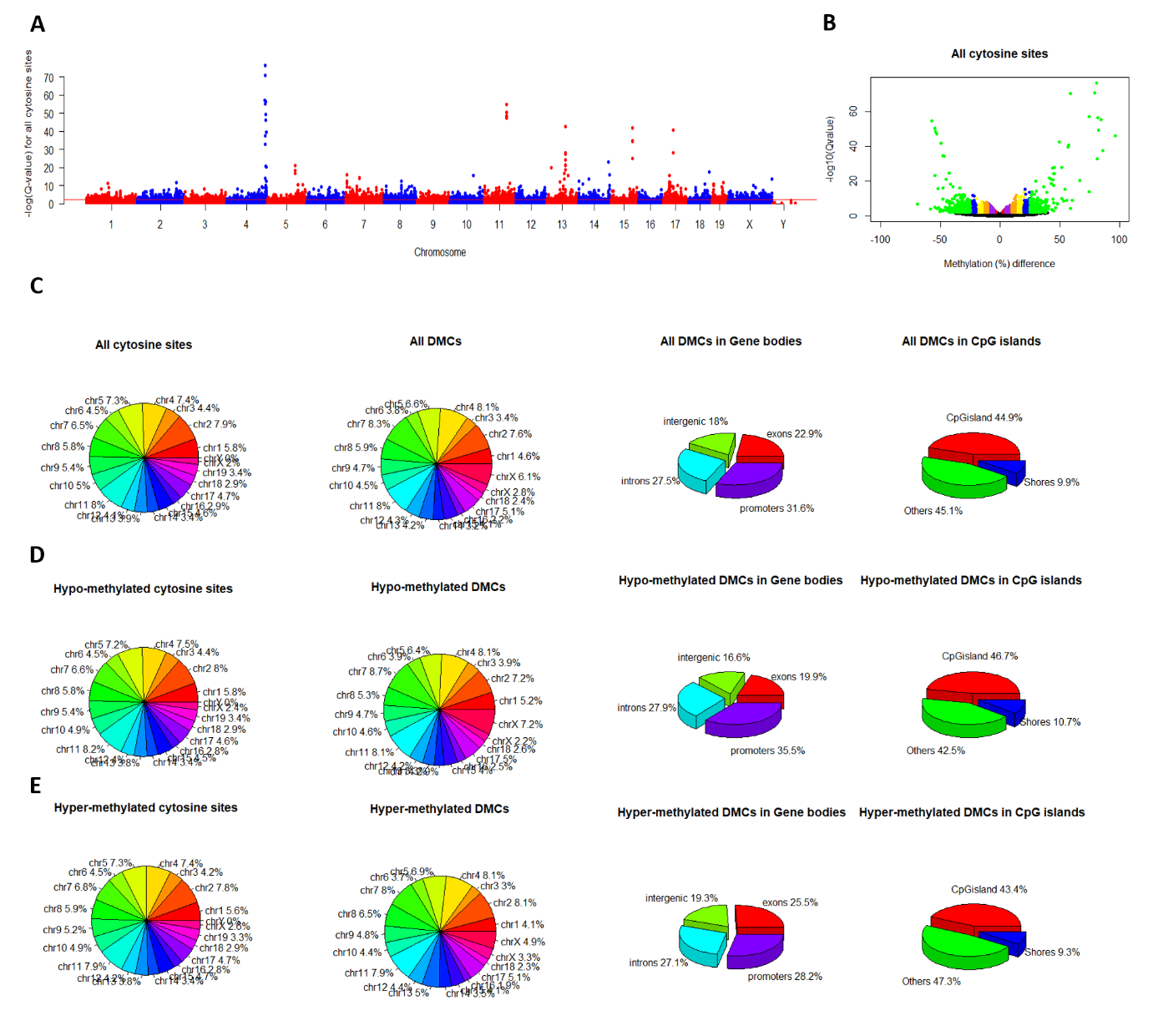

Supplementary figure 2. (**A**) Manhattan plots for all cytosine sites. Note: The red line indicates the significant level of Q-value < 0.01. (**B**) Methylation differences in all cytosine sites. Note: Plots showing red, purple, orange, yellow, blue and green colors indicate genes with a Q-value less than 0.01 and methylation difference (%) greater than 0, 5, 10, 15, 20 and 25, respectively. (**C**), (**D**) and (**E**) Percentages of all, hypo-methylated and hyper-methylated cytosine sites/DMCs in different chromosomes/gene bodies/CpG islands, respectively.

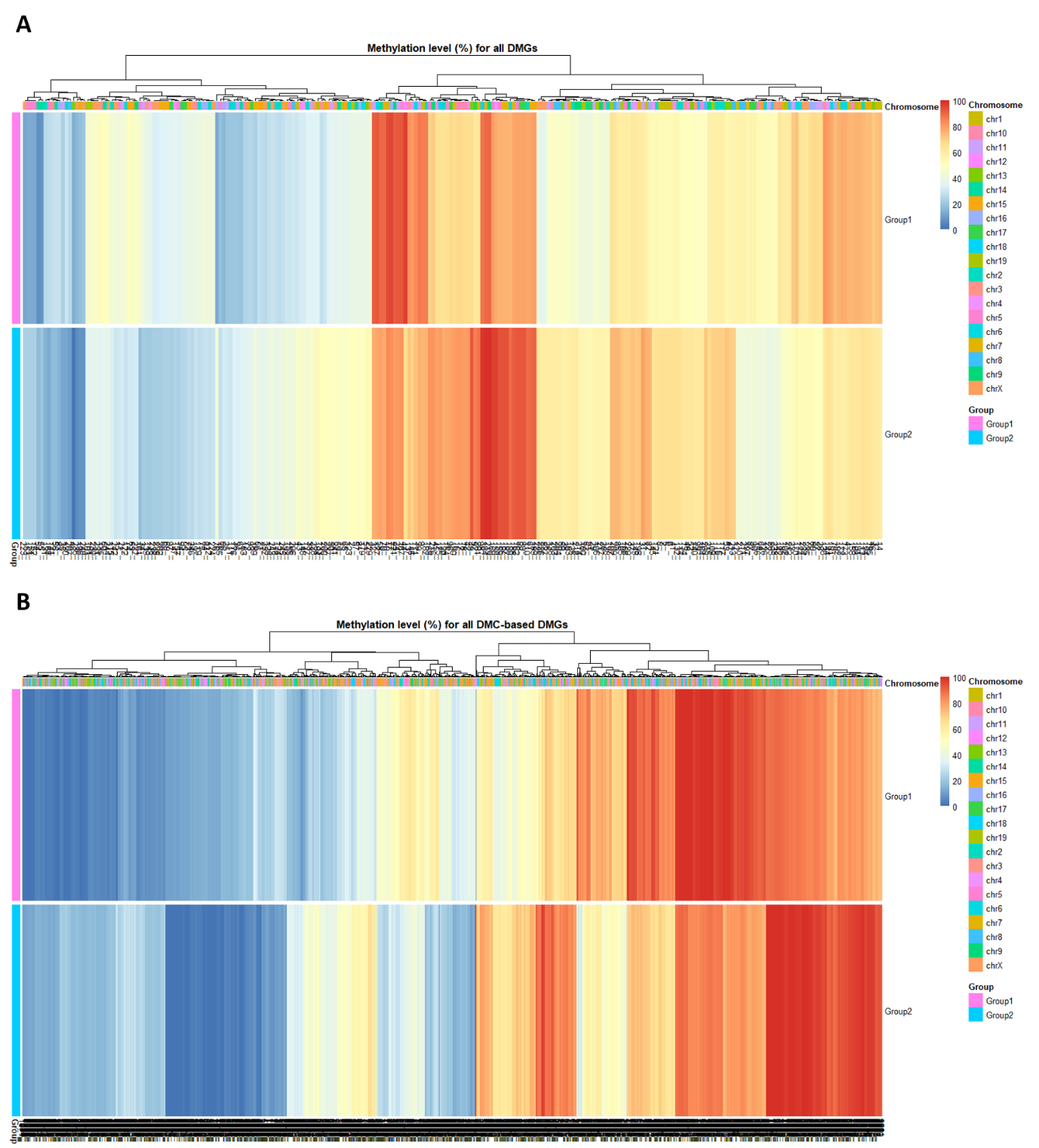

Supplementary figure 3. (**A**) Heat map cluster for methylation levels of all DMGs (n = 246). (**B**) Heat map cluster for methylation levels of all DMC-based DMGs (n = 2022). Note: DMGs and DMC-based DMGs were filter by Significant_filter(qvalue = 0.01, methdiff = 0.1).
